# Seminal root angle is associated with root system architecture in durum wheat

**DOI:** 10.1101/2024.02.27.582216

**Authors:** Yichen Kang, Charlotte Rambla, Shanice V. Haeften, Brendan Fu, Oluwaseun Akinlade, Andries B. Potgieter, Andrew K. Borrell, Emma Mace, David R. Jordan, Samir Alahmad, Lee T. Hickey

**Affiliations:** Centre for Crop Science, The University of Queensland, Queensland Alliance for Agriculture and Food Innovation, Brisbane, QLD, Australia; Salk Institute for Biological Studies, California, USA; School of Agriculture and Food Sciences, The University of Queensland, Brisbane, QLD, Australia; Centre for Crop Science, The University of Queensland, Queensland Alliance for Agriculture and Food Innovation, Gatton, QLD, Australia; Hermitage Research Facility, The University of Queensland, Queensland Alliance for Agriculture and Food Innovation, Warwick, QLD, Australia

**Keywords:** Durum wheat, seminal root angle, root biomass, root system architecture, rhizoboxes

## Abstract

Optimal root system architecture (RSA) is critical for efficient resource capture in soils, hence being an interest in crop breeding. Seminal root angle (SRA) at the seedling stage in durum wheat has been suggested to be a good indicator of RSA. However, research on correlating such lab-based seedling root phenotyping to RSA at later phases of plant growth is limited, resulting in the importance of root trait variation seen in seedlings often being overstated. To explore the role of SRA in modifying RSA at later phases of plant growth, we assessed 11 genotypes contrasting in SRA (wide and narrow), grown in a rhizobox designed for phenotyping root systems of plants during late-tillering. Above-ground traits and root dry mass in different soil depths and across the entire soil volume were measured manually, while root architectural traits were extracted using image analysis and summarised by multiple factor analysis to describe RSA. When comparing the wide and narrow genotypes, no differences were detected for above-ground traits and total root dry mass. However, differences were observed in the allocation of root dry mass at different depths. The wide and narrow genotypes showed distinct RSAs, particularly in the upper soil (0 ‒ 30 cm). The wide genotypes exhibited a ‘spread-out’ root system with dense and thin roots, whereas the narrow genotypes had a compact root system with fewer but thicker roots. Our study demonstrated a clear difference in RSA between the wide and narrow genotypes, highlighting the association between SRA on the direction and distribution of root growth in plants at later growth stages.

## Introduction

Root system architecture (RSA) is the spatial configuration of the root system. RSA influences soil exploration and therefore water and nutrient acquisition, and has been proposed as an important breeding target for a “second green revolution” and the development of climate resilient crops (Lynch, 2007; Ober et al., 2021). The architecture of mature root systems results from a combination of internal (i.e., underlying genetics, developmental influences) and external factors (i.e., environments). The genetic variation in RSA can further respond to environmental cues such as climate, soil depth, as well as its physical and chemical properties, leading to genotype by environment interactions that greatly affect RSA and complicate its selection in breeding programs. Another challenge associated with selection for RSA is the technical difficulties involved in accessing and phenotyping root systems. In particular, it is laborious, costly, and time‒consuming (Johnson, Tingey, Phillips, & Storm, 2001; Trachsel, Kaeppler, Brown, & Lynch, 2011; Wasson et al., 2014) to measure roots under field conditions, despite the limitations of the data they produce.

In contrast, root phenotyping in controlled environments can help address some of the above-mentioned bottlenecks, enabling more reliable phenotypic data to be obtained and permitting measurement at a relatively high throughput. To date, there are number of custom phenotyping platforms developed that allow the study of root systems (Nagel et al., 2012; Figueroa-Bustos, Palta, Chen, & Siddique, 2018). These root phenotyping platforms consist of root growth chambers and digital imaging of entire root systems. Recent advances in image processing software have allowed users to extract an array of traits that describe root architecture within the root growth chambers. Thus, the combination of root observation systems and new developments in computer‒assisted electronic image analysis presents an opportunity to enhance the capacity for phenotyping RSA in root growth chambers, with the intention that these traits are relevant to the field performance of the genotypes.

RSA is typically studied by analysing the sub-root system component traits, e.g., length, number, and angle of roots (Rich & Watt, 2013). The measurement of root component traits or the utilisation of molecular markers linked to these traits could provide an indirect selection method for genotypes with desirable RSAs. One root trait of particular interest is the angle of seminal roots. In durum wheat (*Triticum durum* Desf.), previous studies suggest that seminal root angle (SRA) at the seedling stage could be a useful ‘proxy’ for the root angle of mature root systems (Maccaferri et al., 2016; Alahmad et al., 2019). SRA affects the direction of root growth during early developmental stages and therefore has the potential to influence the later horizontal and vertical exploration of the soil profile. In bread wheat, genetic loci for stay‒ green, an integrated drought adaptation trait, have been found to co‒locate with loci for root angle (Christopher et al., 2018). Similarly, previous studies in sorghum have also reported co‒ locations of genetic loci for stay‒green traits and root angle (Mace et al., 2012; Borrell et al., 2014). Since one of the important mechanisms underlying stay‒green is associated with increased post‒flowering water uptake in terminal drought environments, a root system with a steep angle should result in a deep rooting system that enhances access to deep soil water (Manschadi, Christopher, Devoil, & Hammer, 2006; Singh, van Oosterom, Jordan, & Hammer, 2012; Borrell et al., 2014; Borrell et al., 2022). Therefore, SRA may be a major determinant of the mature RSA, making it a promising candidate for selection that can be conducted in the early growth stages to accelerate the breeding process.

SRA of seedling plants can be easily measured using lab- or glasshouse-based phenotypic screening protocols (Cane et al., 2014; Richard et al., 2015), which are amenable to large numbers of genotypes and have contributed to the identification of genetic loci for SRA in wheat (Richard et al., 2015), barley (Robinson et al., 2016), sorghum (Singh et al., 2012; Menamo et al., 2023) and durum wheat (Cane et al., 2014; Maccaferri et al., 2016; Alahmad et al., 2019; Alemu et al., 2021). Although this shows the potential of using SRA to enable high-throughput phenotyping and indirect selection for optimal root systems, it remains unclear to what extent SRA influences the RSA. Previously, significant genotypic variations in SRA have been demonstrated in a durum wheat nested association mapping (NAM) population (Alahmad et al., 2019). Based on this, the present study aimed to elucidate the impact of SRA on RSA at later growth stages. Here, we conducted a short-term glasshouse experiment extended to late-tillering stage to quantify the RSA for two distinct groups of durum NAM lines categorised according to contrasting SRA.

## Materials and methods

### Plant material

Previously, a durum NAM population was developed by crossing eight ICARDA ‘founder’ lines with two Australian durum ‘reference’ varieties Jandaroi and DBA Aurora (Alahmad et al., 2022). For this study, 11 genotypes including the reference parent DBA Aurora and 10 NAM lines, based on the SRA screenings by Alahmad et al. (2019), were selected for divergent SRAs (wide and narrow) (Table 1).

**Table 1.** The pedigree information, seminal root angle (SRA) group and value of the genotypes surveyed in this study.

| Genotype | Pedigree | Group | SRA value<sup>1</sup> (°) |
| --- | --- | --- | --- |
| DBA Aurora | Tamaroi*2/Kalka//RH920318/Kalka//Kalka*2/Tamaroi | Wide | 81.1 |
| 1_17 | DBA Aurora/Fastoz7 | Wide | 86.3 |
| 1_42 | DBA Aurora/Fastoz7 | Wide | 79.3 |
| 1_63 | DBA Aurora/Fastoz7 | Wide | 82.6 |
| 1_99 | DBA Aurora/Fastoz7 | Wide | 80.4 |
| 1_107 | DBA Aurora/Fastoz7 | Wide | 71.4 |
| 3_51 | DBA Aurora/Fastoz8 | Narrow | 51.9 |
| 3_86 | DBA Aurora/Fastoz8 | Narrow | 52.8 |
| 3_88 | DBA Aurora/Fastoz8 | Narrow | 38.5 |
| 6_17 | Jandaroi/Fastoz8 | Narrow | 41.5 |
| 6_21 | Jandaroi/Fastoz8 | Narrow | 50.9 |

<sup>1</sup> Values of SRA were the genotype means for DBA Aurora and the NAM lines obtained from the association mapping study by Alahmad et al. (2019). The SRA, which is the angle between the first pair of seminal roots, was measured at five days after sowing using the clear pot method (Richard et al., 2015; Alahmad et al., 2019).

### Glasshouse rhizobox experiment

A glasshouse experiment was conducted in October 2020 in a temperature‒controlled glasshouse (17 ± 2°C) under natural light conditions at The University of Queensland, Brisbane,Australia (27.50°S, 153.01°E). Four‒sided open-top rhizoboxes were constructed from timber, with each rhizobox measuring 90 cm tall × 40 cm wide. The bottom of each rhizobox was sealed with water-permeable shade‒cloth to hold the soil. The rhizoboxes were placed upright in rectangular trays, and braced across the top edges with a wooden stay (Figure 1A), and filled with UQ23 potting mix (70% composted pine bark 0 ‒ 5mm, 30% cocoa peat, mineral fertiliser) pre‒mixed with 2 g/L of Osmocote slow‒release fertiliser. All rhizoboxes were thoroughly watered before sowing.

**Figure 1.**
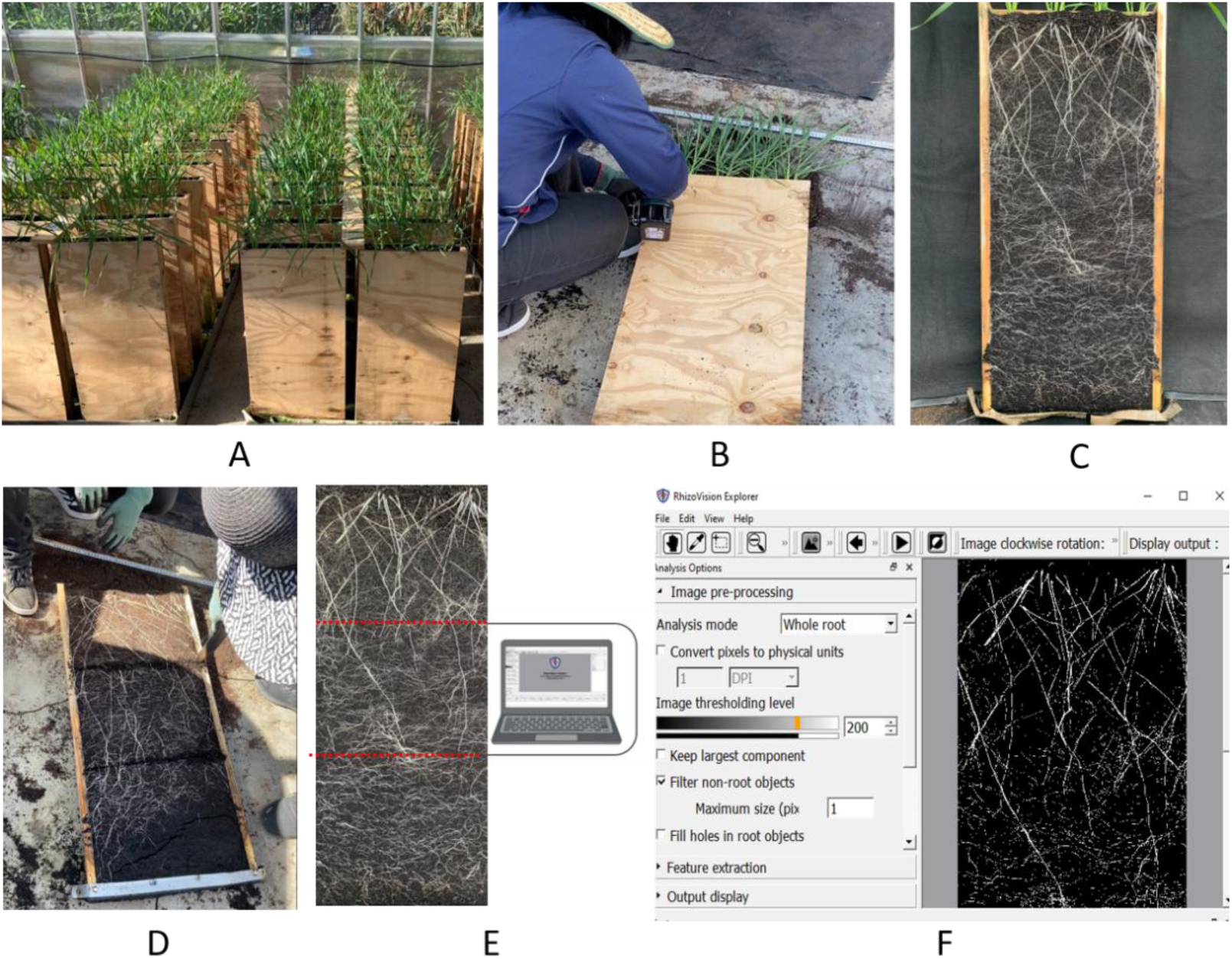
The workflow of the rhizobox experiment for root phenotyping. (A) Seedlings were grown for six weeks in rhizoboxes in a temperature‒controlled glasshouse. (B) Opening of the rhizoboxes at the end of the experiment. (C) Imaging using a smartphone. (D) Separating the soil profile into three equal sections (0 ‒ 30, 30 ‒ 60, and 60 ‒ 90 cm) for root washing. (E) Cropping images into three equal sections for analysing root system architecture (RSA) in differing soil depths. (F) Working photo of image segmentation and analysis with RhizoVision Explorer software.

At sowing (7/10/2020), to avoid confounding genetic effects, uniformly sized seeds were selected closest to the median size for each genotype. Eight seeds of a single genotype were sown embryo down at a depth of 3 cm, with two seeds sown in each of the four holes that were evenly distributed in each rhizobox. Plants were thinned to four homogenous and healthy seedlings after germination. Rhizoboxes were arranged in a randomised complete block design, with six rhizoboxes (replicates) of each genotype. Rhizoboxes were watered regularly to ensure plants were supplied with non‒limiting water throughout the experiment. For all plants, nutrients and disease control measures were applied as necessary.

To determine when to open the rhizoboxes, (since the experimental rhizoboxes were kept enclosed until the end of the experiment), we created a pilot rhizobox planted with DBA Aurora seeds and opened it once a week to monitor root growth. The pilot rhizobox was subjected to the same sowing date and growing conditions as the experimental rhizoboxes.

### Root imaging

The experiment ceased six weeks after sowing (late-tillering) when roots in the pilot rhizobox had almost reached the bottom of the wooden frame. All experimental rhizoboxes were opened for root imaging using a high‒quality smartphone camera (Apple iPhone 11) (Figure 1B, C), and plants were harvested. When taking images, each rhizobox was carefully straightened and its four edges were aligned parallel to the photo frame of the camera. These images were then cropped to retain only the soil section comprising the whole root system. To further explore the RSA in different soil depths, namely the 0 ‒ 30, 30 ‒ 60, and 60 – 90 cm sections of the rhizobox, we used the online image cropping tool at https://www.imgonline.com.ua/eng/ to cut the cropped images of the intact root system (0 – 90 cm) into three images of equal size (Figure 1E). Consequently, each experimental rhizobox had four images that showed the whole root system (0 – 90 cm), and the root systems in the 0 – 30 cm, 30 – 60 cm and 60 – 90 cm soil depths.

All images were processed using RhizoVision Explorer version 2.0.3 software (Seethepalli et al., 2021) (Figure 1F), selecting the ‘Whole root’ analysis mode, which unlike the ‘Broken root’ mode, gives additional information on the growth direction of the root system (e.g., average root orientation, shallow angle frequency and steep angle frequency), and filtering non‒root objects with default size value and enabling pixel to millimetre conversion for statistical analysis. The diameter ranges 1, 2 and 3 were set as default, and corresponded to 0 – 2 mm, 2 – 5 mm and above 5 mm, respectively. A total of 32 root imaging traits were extracted from RhizoVision Explorer, which provided detailed information on the RSA (Table 2).

**Table 2.** Traits obtained from the rhizobox experiment. Root imaging traits analysed with RhizoVision Explorer were further classified into different root trait groups. Abbreviations and root trait groups of root imaging traits followed Adu et al. (2022), description of root imaging traits can be found in Seethepalli et al. (2021).

| Trait | Abbreviations | Root trait group | Unit |
| --- | --- | --- | --- |
| Seminal root angle | SRA | — | ° |
| Average root orientation | ARO | Angle | ° |
| Network area | NeA | Area | mm<sup>2</sup> |
| Convex area | CoA | Area | mm<sup>2</sup> |
| Solidity | — | Area | — |
| Lower root area | LRA | Area | mm<sup>2</sup> |
| Surface area | SA | Area | mm<sup>2</sup> |
| Average hole size | AHS | Area | mm<sup>2</sup> |
| Projected area diameter—range 1 | PAD1 | Area | mm<sup>2</sup> |
| Projected area diameter—range 2 | PAD2 | Area | mm<sup>2</sup> |
| Projected area diameter—range 3 | PAD3 | Area | mm<sup>2</sup> |
| Surface area diameter—range 1 | SAD1 | Area | mm<sup>2</sup> |
| Surface area diameter—range 2 | SAD2 | Area | mm<sup>2</sup> |
| Surface area diameter—range 3 | SAD3 | Area | mm<sup>2</sup> |
| Average diameter | AvD | Diameter | mm |
| Median diameter | MeD | Diameter | mm |
| Maximum diameter | MxD | Diameter | mm |
| Total root length | TRL | Length | mm |
| Perimeter | — | Length | mm |
| Root length diameter—range 1 | RLD1 | Length | mm |
| Root length diameter—range 2 | RLD2 | Length | mm |
| Root length diameter—range 3 | RLD3 | Length | mm |
| Holes | — | Number | — |
| Shallow angle frequency | ShAF | Number | — |
| Medium angle frequency | MAF | Number | — |
| Steep angle frequency | StAF | Number | — |
| Median number of roots | MNR | Number | — |
| Maximum number of roots | MxNR | Number | — |
| Number of root tips | NRT | Number | — |
| Volume | — | Volume | mm<sup>3</sup> |
| Volume diameter—range 1 | VDR1 | Volume | mm<sup>3</sup> |
| Volume diameter—range 2 | VDR2 | Volume | mm<sup>3</sup> |
| Volume diameter—range 3 | VDR3 | Volume | mm<sup>3</sup> |
| Root dry mass | — | — | mg |
| Stem number | — | — | — |
| Leaf area | — | — | mm<sup>2</sup> |
| Shoot dry mass | — | — | g |
| Root-to-shoot ratio | — | — | — |

### Root and shoot manual measurements

At the end of the experiment, shoots and roots were harvested. To determine root distribution with soil depth, root subsamples were cut from the root system into 0 ‒ 30, 30 ‒ 60, and 60 ‒ 90 cm sections and roots from each section were washed clean (Figure 1D). Considering that root washing is a time‒consuming process, only two replicates of each genotype were subjected to root washing, which provided a total of 12 replicates for the wide SRA and 10 replicates for the narrow SRA for analysis. Shoot samples from all rhizoboxes were carefully separated into leaves and stems for measuring total leaf area using a LICOR 3100 leaf area meter and counting stem number before drying and weighing. Shoot and root samples were dried at 65 °C in an air‒forced oven for 72 h to obtain the total shoot dry mass and root dry mass of each section. The root‒to‒shoot ratio was calculated as total root dry mass/(total shoot dry mass x 1000). All trait values were analysed on a four‒plant basis (rhizobox-level).

### Statistical analysis

Data were analysed and visualised in the R statistical environment (R Core Team, 2013). To summarise 32 root imaging traits obtained from RhizoVision Explorer, we structured these traits into six different root trait groups as described by Adu et al. (2022), except for surface area with different diameter ranges (i.e., 1, 2 and 3), which was grouped to root area in our study. The list of root imaging traits and their corresponding root trait groups (i.e., root angle, area, diameter, length, number and volume) can be found in Table 2. To interpret RSA, we performed multiple factor analysis to assess the correlative structure of the groups of root imaging traits measured on the intact root system (0 – 90 cm) and on the roots in different soil depths (0 ‒ 30, 30 ‒ 60, and 60 – 90 cm). Briefly, multiple factor analysis computed the principal component analysis (PCA) twice, with the first PCA conducted for each group of root imaging traits, then the resulting normalised group-based PCA data were used to compute the second PCA (global PCA) to assess the relationships between root trait groups. The group similarity that reflected the correlation between two groups was evaluated by the RV coefficient (ranging from 0 – 1, the higher the RV, the stronger the relationship), a multivariate generalisation of the *squared* Pearson correlation coefficient (Abdi, Williams, & Valentin, 2013). The R package “FactoMineR” was used to perform multiple factor analysis (Le, Josse, & Husson, 2008).

For the comparison of RSA, we used the axis scores of the first two principal components (PCs) extracted from each multiple factor analysis. To gain a better understanding of the PCs, pair-wise Pearson’s correlation was conducted between the root imaging traits and the scores of the first two PCs of each multiple factor analysis. Only the root imaging traits that showed a significant correlation with the PC scores were retained to explain the corresponding PC. To make the analysis more manageable, we only interpreted the PCs that were significantly different between the wide and narrow SRAs to demonstrate the difference in RSA.

For shoot traits, overall root dry mass (0 ‒ 90 cm), overall RSA PCs (0 ‒ 90 cm) and root‒to‒ shoot ratio, significance testing of the genotype or SRA effects was performed via one-way analysis of variance (ANOVA). Whereas for root dry mass and RSA PCs assessed at various depths (0 ‒ 30, 30 ‒ 60, and 60 – 90 cm), two-way ANOVA was used to identify the effects of soil depth and genotype or SRA. The residual normal distribution and homoscedasticity of all models were ascertained by Shapiro’s test and plotting residuals against quantiles. A *t*-test or post-hoc test was followed up to compare the arithmetic means of the wide and narrow SRA genotypes. Unless otherwise stated, statistical significance was at *P* < 0.05.

## Results

### The relationship between seminal root angle and distribution of root biomass

The phenotypic variation differed among traits measured manually in the rhizobox experiment, with the lowest being shoot dry mass (CV = 20.4%) and the highest being the allocation of root dry mass within the 60 ‒ 90 cm soil section (CV = 47.8%) (Table 3). The results of the ANOVA suggested that genotype was a significant factor affecting most of these traits. Between the wide and narrow genotypes, differences were observed for root dry mass in the 0 ‒ 30 cm soil section (*P* < 0.01). Specifically, in this section, wide genotypes had significantly increased root dry mass compared with the narrow genotypes (narrow = 932 mg; wide = 1258 mg) (Figure 2, Table 3).

**Figure 2.**
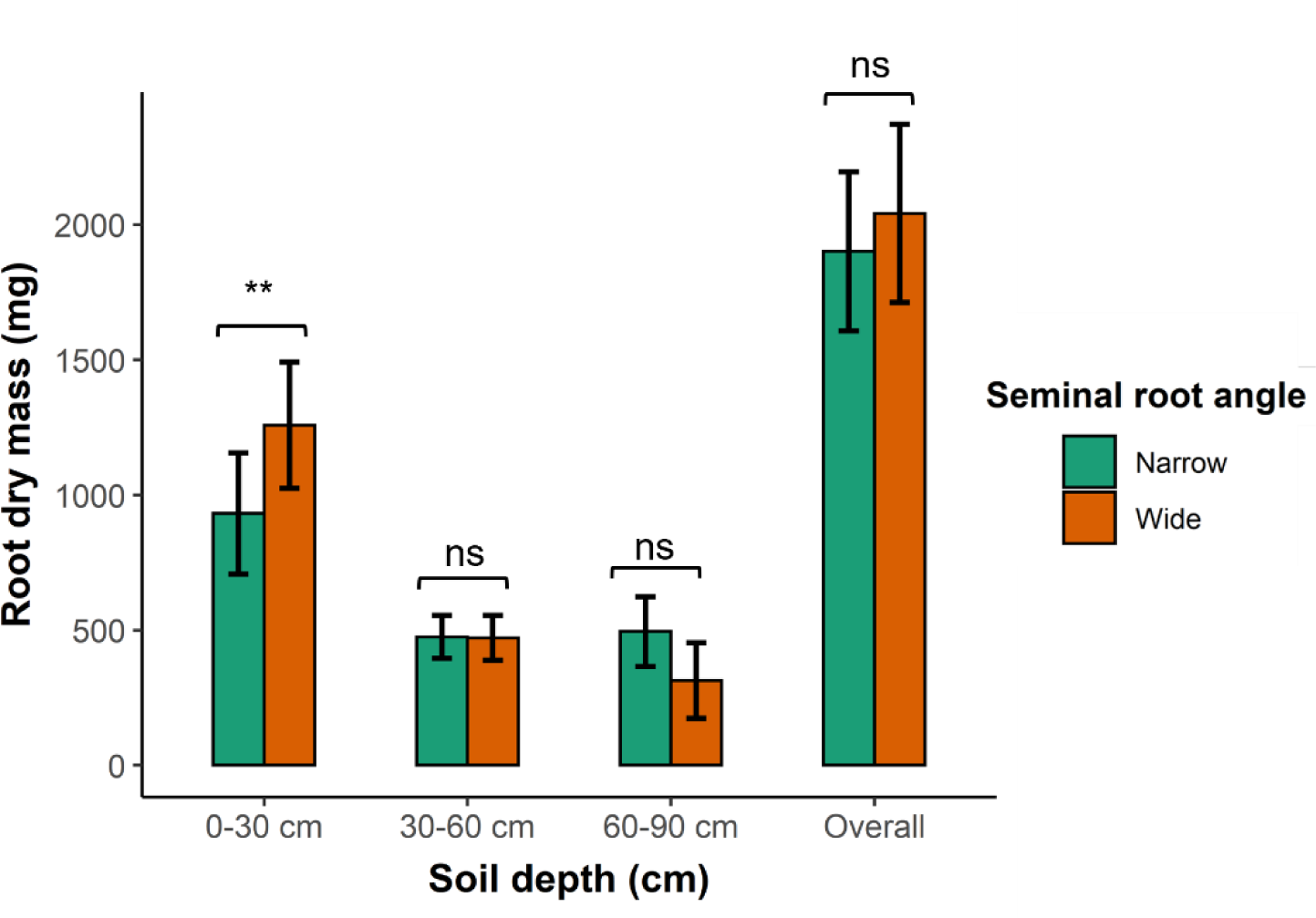
Root dry mass in different soil depths of rhizoboxes for groups of genotypes contrasting for seminal root angles (i.e., wide and narrow). The comparison between the wide and narrow genotypes indicates that the means are non‒significant (ns), or significant at ** *p* < 0.01, respectively. Error bars represent the 95% CIs for means. The statistics can be found in Table 3.

**Table 3.** Descriptive statistics for traits manually measured in the rhizobox experiment, including mean, range and coefficients of variation (CV), and the significance of the genotype main effect and comparison between the wide and narrow SRAs.

| Trait | Depth | Descriptive statistics |  |  | Genotype effect<br>(*P*-value) | Mean |  | Comparison<br>(*P*-value) |
| --- | --- | --- | --- | --- | --- | --- | --- | --- |
|  |  | Mean | Range | CV (%) |  | Wide | Narrow |  |
| Root dry mass (mg) | 0 – 30 cm | 1109 | 618 – 2451 | 37.8 | < 2.49E – 07 | 1258 | 932 | < 0.01 |
|  | 30 – 60 cm | 473 | 296 – 668 | 20.5 |  | 471 | 475 | 1.00 |
|  | 60 – 90 cm | 396 | 113 – 857 | 47.8 |  | 314 | 495 | 0.35 |
| Overall root dry mass (mg) | - | 1978 | 1125 – 3075 | 24.5 | < 0.01 | 2042 | 1901 | 0.51 |
| Stem number | - | 7 | 3 – 11 | 27.7 | 1.30E – 06 | 7 | 6 | 0.38 |
| Leaf area (mm<sup>2</sup>) | - | 424 | 225 – 669 | 23.7 | < 0.001 | 400 | 431 | 0.67 |
| Shoot dry mass (g) | - | 2.33 | 1.30 – 3.48 | 20.4 | < 0.05 | 2.39 | 2.23 | 0.32 |
| Root-to-shoot ratio | - | 0.86 | 0.57 – 1.30 | 23.6 | 0.09 | 0.87 | 0.85 | 0.88 |

### Summarising root system architecture by multiple factor analysis

The multiple factor analysis was conducted on the intact root system (0 – 90 cm) and on the roots in different soil depths (0 ‒ 30, 30 ‒ 60, and 60 – 90 cm) (Table 4). The results suggested that 77.3 ‒ 82.5% of the total variation in RSA was captured by PC1 and PC2, with 63.5 ‒ 69.8% and 9.9 ‒ 17.8% explained by PC1 and PC2, respectively. Since PC3 accounted for the least variation in RSA (it only contributed 5.4 ‒ 7.7%), it was excluded from further analyses. Among six root trait groups, root angle had the least contribution to PC1, ranging between 9.4 – 12.4% (Figure 3 and Supplemental table S1). Concerning PC2, it was primarily associated with root angle and/or root area, where the contribution of root angle was as high as 49%. Moreover, the relationship between root trait groups showed that except for root angle, the other five groups were highly related to each other (RV = 0.68 – 0.92) (Table 5). Root angle showed less of a relationship with other groups (RV = 0.21 – 0.67), with the strongest correlation being observed with root number, with RV coefficients ranging from 0.45 to 0.67.

**Figure 3.**
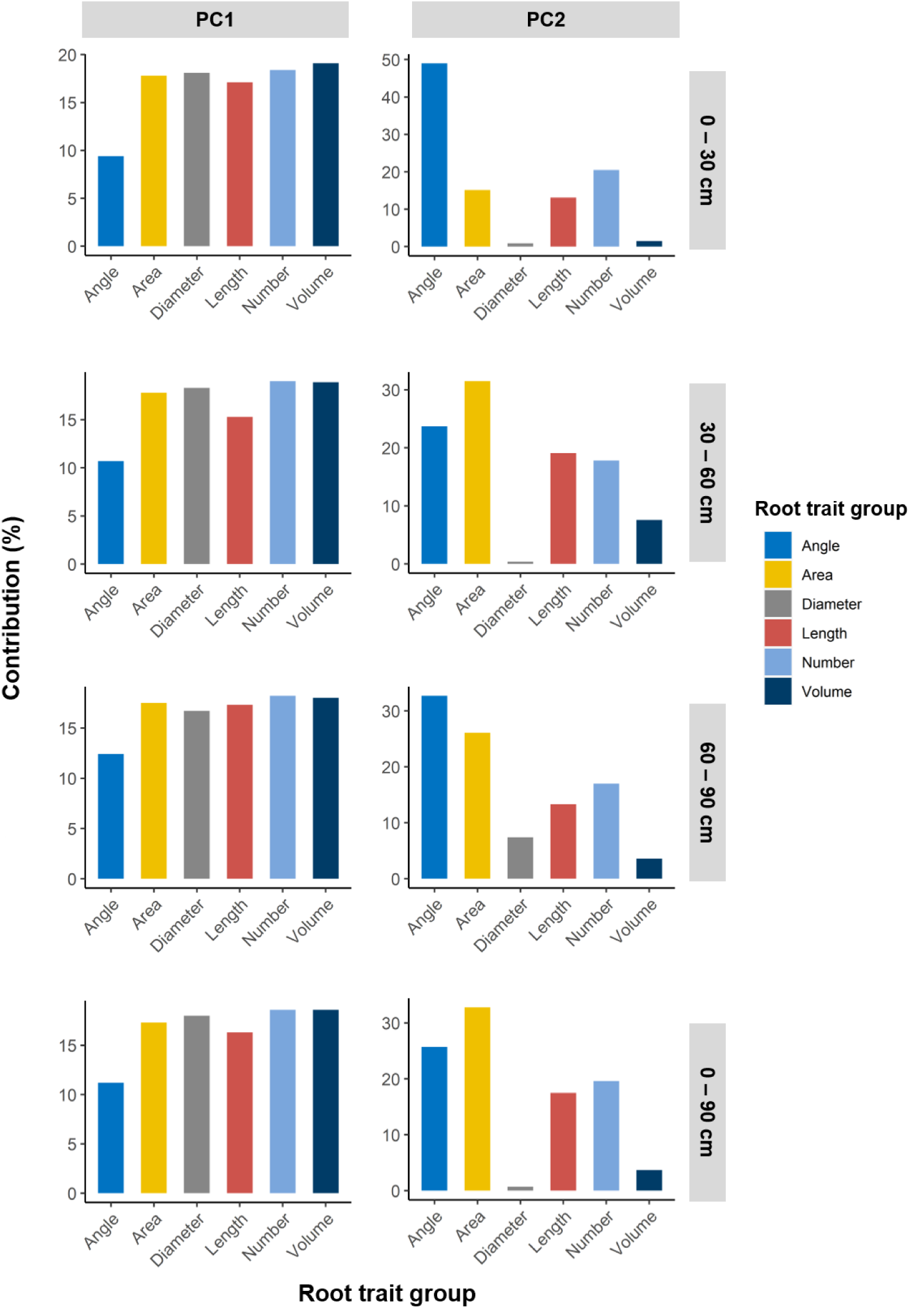
Barplots displaying the contribution of root trait groups to the first two principal components (PC1 and PC2) of each multiple factor analysis.

**Table 4.** Contribution of the first three principal components (PC1, PC2 and PC3) of the multiple factor analysis conducted on the intact root system (0 – 90 cm) as well as on the roots in different soil depths (0 ‒ 30, 30 ‒ 60, and 60 – 90 cm).

| Multiple factor analysis | Contribution (%) |  |  |  |
| --- | --- | --- | --- | --- |
|  | PC1 | PC2 | PC3 | PC1 + PC2 |
| 0 – 30 cm | 68.3 | 12.1 | 6.9 | 80.4 |
| 30 – 60 cm | 64.7 | 17.8 | 5.4 | 82.5 |
| 60 – 90 cm | 69.8 | 9.9 | 5.9 | 79.7 |
| Overall (0 – 90 cm) | 63.5 | 13.7 | 7.7 | 77.3 |

**Table 5.**
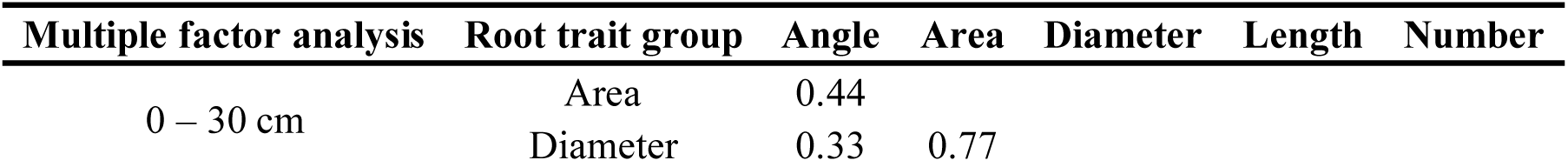
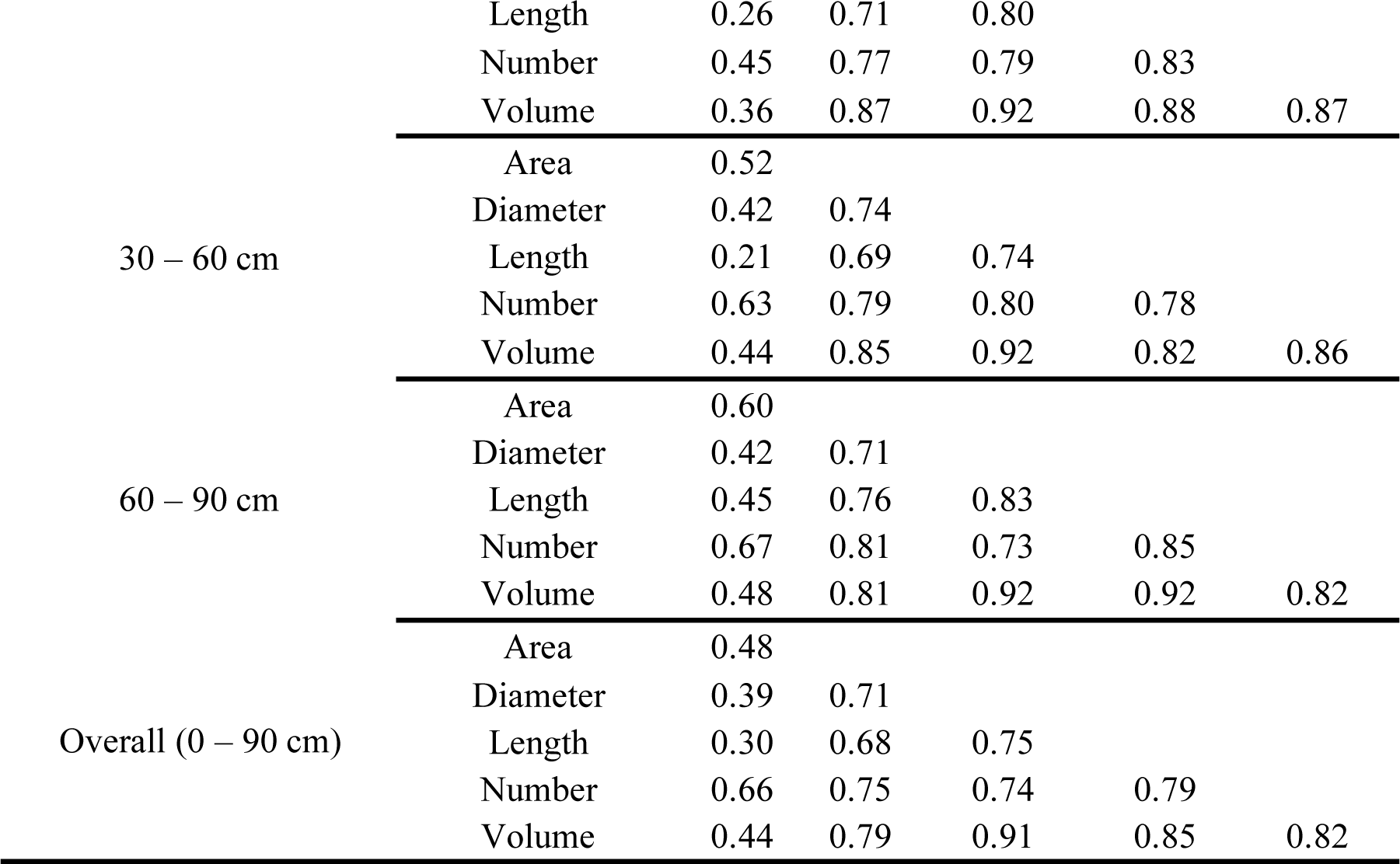
Pair‒wise RV coefficients from each multiple factor analysis performed on six root trait groups.

### Seminal root angle influenced the proportion of roots in the top 0-30 cm soil section

The genotype means for the scores of the first two PCs are presented in Supplemental Table S2. The rankings of these 11 genotypes for PC1 and PC2 provide an approximate indication of the relationship between SRA and RSA. The results showed a good separation between the wide and narrow genotypes for PC1 of the multiple factor analysis conducted on the roots in the 0 ‒ 90 cm and 0 ‒ 30 cm depths, where the wide and narrow genotypes were additionally well separated for PC2 in the 0 ‒ 30 cm section (Figure 4). In contrast, no obvious trend of difference between the wide and narrow genotypes was observed for PCs at other depths (i.e., 30 – 60 cm and 60 – 90 cm). These results indicated the potential effectiveness of SRA in creating root systems with divergent RSA, which was likely driven by the difference in RSA in the 0 ‒ 30 cm section.

**Figure 4.**
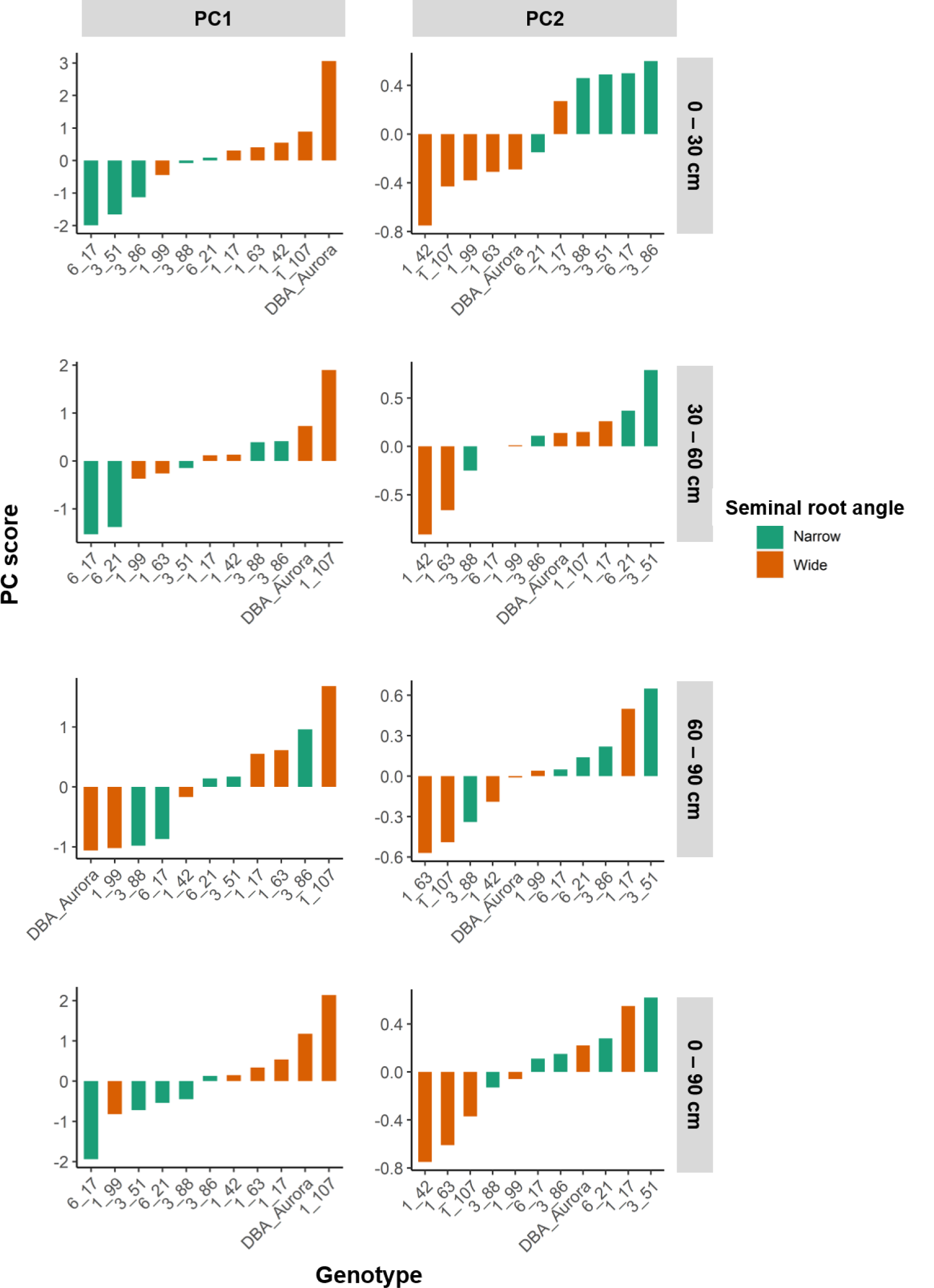
Rankings of durum wheat genotypes for PC1 and PC2 of each multiple factor analysis.

To further investigate the association between SRA and RSA in the soil, we compared the wide and narrow genotypes for the first two PCs in different soil depths as well as across the entire soil profile (Figure 5). When assessed across the entire soil profile (0 ‒ 90 cm), PC1 was significantly different between the wide and narrow genotypes (*P* < 0.05). When assessed in different soil depths, the wide and narrow genotypes differed significantly in the 0 ‒ 30 cm soil section (*P* < 0.01) for both PC1 and PC2. These results confirmed that the different overall RSA between the wide and narrow genotypes was mainly due to their difference in RSA in the 0 ‒ 30 cm section. Specifically, in this section, the narrow genotypes displayed lower PC1 than the wide genotypes but higher PC2 than the wide genotypes.

**Figure 5.**
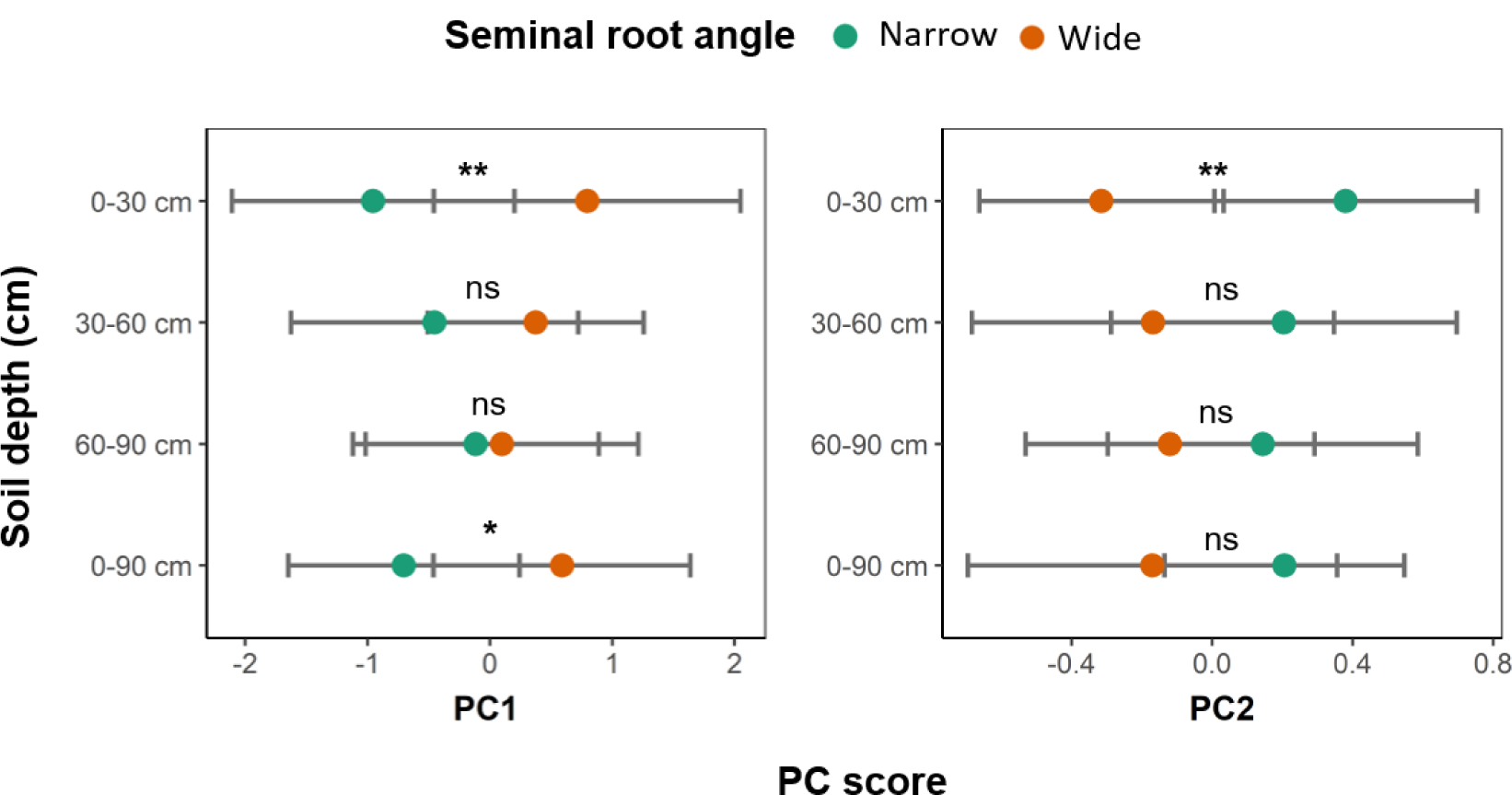
Comparison of the PC1 and PC2 between the wide and narrow SRAs in the soil profile. Statistically significant differences are indicated by asterisks, where *, ** and ns denote significance at *p* < 0.05 and *p* < 0.01 and non‒significance, respectively. Results are displayed as means with 95% CIs, with details provided in the Supplemental table S3.

### Wide and narrow genotypes differed in root growth direction and morphology

Since the RSAs of wide and narrow genotypes were differentiated by PC1 and PC2 in the 0 ‒ 30 cm soil section, and by PC1 in the 0 ‒ 90 cm soil section, we focused on the interpretation of these PCs. To do this, correlations were analysed between individual root imaging traits and each of the PC scores. Traits showing significant correlation with PC are detailed in Supplemental Table S4.

For PC1 in the 0 ‒ 30 cm section, its higher values were correlated with smaller average root orientation (i.e., roots grew in a more horizontal direction, as root orientation is measured in relation to the horizontal line), larger projected and surface areas with thinner roots (diameter ranges 1 and 2), smaller projected and surface areas with thicker roots (diameter range 3), and an overall smaller root area (i.e., SA, NeA and CoA). Higher PC1 values were also correlated with thinner roots, greater total root length, greater number of roots and greater number of roots with shallow angle, higher volume with thin roots (diameter ranges 1 and 2), smaller volume with thick roots (diameter range 3), and an overall smaller root volume. For PC1 in the 0 ‒ 90 cm section, generally this showed the same correlation trend as in the 0 ‒ 30 cm section. This again confirmed that SRA primarily influenced RSA in the 0 ‒ 30 cm section.

For PC2 in the 0 ‒ 30 cm soil section, its higher values were correlated with greater average root orientation (so steeper root growth), larger root area with thick roots (diameter range 3) and an overall larger root area, as well as greater root length with thick roots (diameter range 3) and a greater total root length, a greater number of roots with steep angle, and a smaller number of roots with shallow angle. However as PC2 was dominated by root angle (contribution = 49%) and root number (contribution = 20.5%) in the 0 ‒ 30 cm section (Supplemental Table S1), its interpretation was therefore simplified by referencing its correlation with root angle and number of groups alone. As such, higher PC2 values were correlated with steeper root growth, greater number of roots with steep angle and smaller number of roots with shallow angle, all features indicating a steeper root system.

To conclude, the higher PC1 and lower PC2 of the wide genotypes in the 0 ‒ 30 cm section indicated a horizontal root distribution with a greater total root length, a larger number, surface area and volume of thin roots (Figure 6). Conversely, the narrow genotypes had a more vertical root distribution with a shorter total root length, but a larger number, surface area and volume of thick roots.

**Figure 6.**
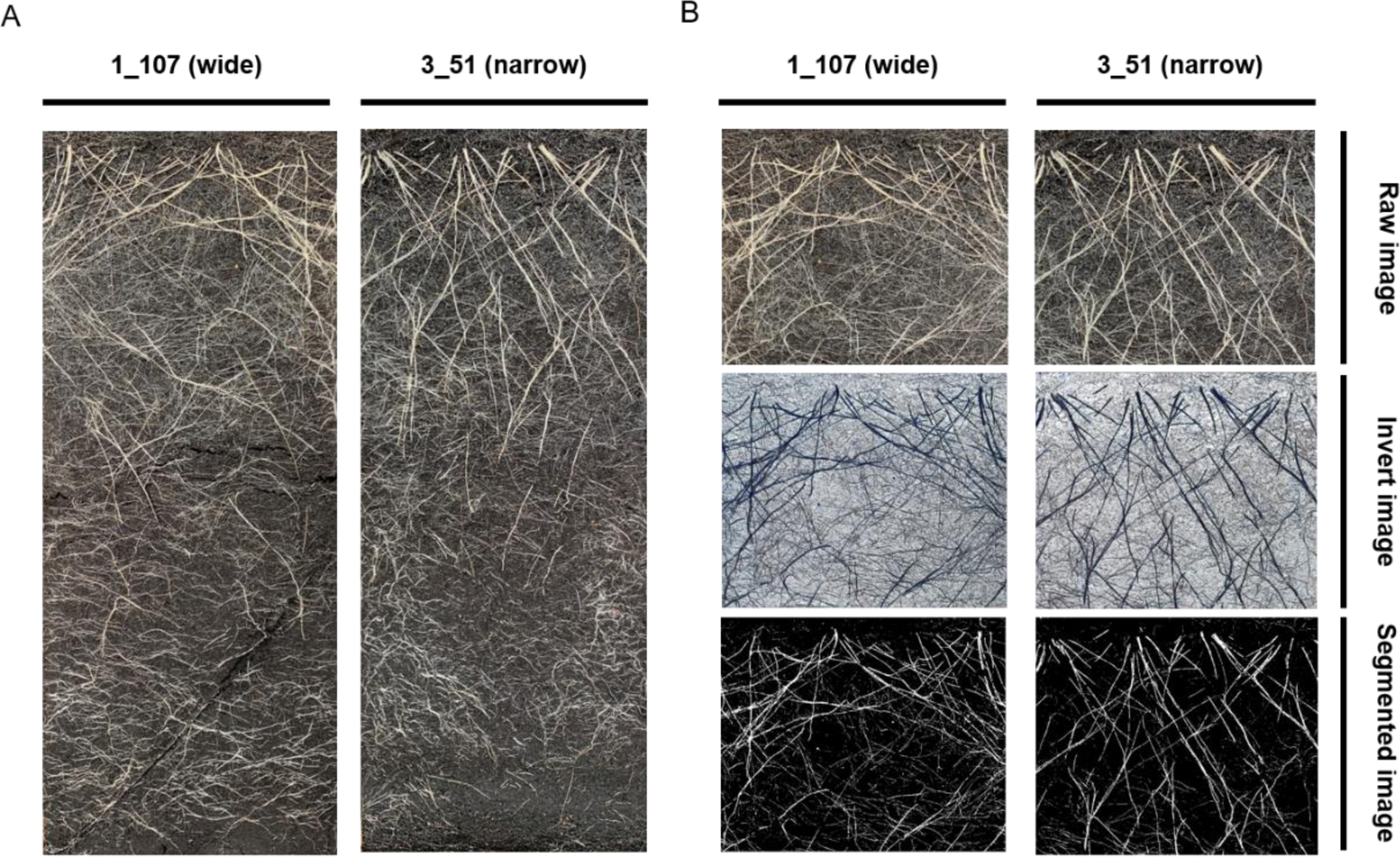
(A) Representative root images of two genotypes with contrasting seminal root angles, (B) and visualisation of roots in the 0 ‒ 30 cm section in RhizoVision Explorer.

## Discussion

This study investigated the association between seedling root angle (SRA) and different aspects of root system architecture (RSA) measured in later-tillering plants, with the aim of providing evidence that low‒cost high‒throughput SRA phenotyping methods (e.g., ‘clear pot’ method) can be used to make inferences about subsequent growth of the root system in older plants. Although the use of rhizoboxes does not establish a direct connection between SRA and the fully developed root system or production at the field level, it represents a good compromise between time efficiency and handling capacity for studying root systems at intermediate time points. Understanding root development throughout various growth stages is crucial, and the earlier RSA may significantly influence the mature RSA. Coupled with imaging analysis, quantitative descriptions of RSA facilitated complex comparisons between the narrow and wide SRA-associated root systems. The observed differences in RSA of genotypes contrasting for SRA indicate that the high-throughput SRA phenotyping may be a useful tool in predicting RSA expressed at later growth stages, which could be applied in durum wheat breeding programs to develop optimal root systems for specific environments.

### Seminal root angle may regulate root growth direction over time independent of total root biomass

Root biomass is an important indicator of the size of a root system (Adiku, Ozier-Lafontaine, & Bajazet, 2001). A larger root system can improve the uptake of water and nutrients and hence increase yield performance. However, the benefit of a large root system is very context dependent, relying greatly on the differences in the depth at which soil resources are present under variable growing conditions (Palta et al., 2011). In some circumstances, for example, in the presence of terminal drought, early exploitation of soil water deep in the profile may lead to early depletion of soil water and limited growth of yield components, ultimately reducing grain yield (Figueroa-Bustos, Palta, Chen, Stefanova, & Siddique, 2020). In fact, the ‘Green Revolution’ resulted in the unintentional selection for smaller root systems in modern wheat cultivars (Waines & Ehdaie, 2007). Thus, several decades of breeding for yield has progressively reduced root biomass, root length and root length density in modern cultivars, while nitrogen uptake has increased (Aziz, Palta, Siddique, & Sadras, 2017; Zhang, Du, & Li, 2020). This highlights the importance of an efficient root system rather than simply a large one (van Oosterom et al., 2016). Thus, root traits that can improve the efficiency of the root system in acquiring soil resources would be better targets for selection than root size *per se* (Lynch, 2007).

In the rhizobox experiment, the wide and narrow genotypes did not differ in total root biomass, but did show significant variation in the distribution of root biomass across the soil profile. This suggests the potential for SRA to change root biomass distribution without influencing the total root biomass. Thus, selection for SRA may not result in a carbon trade‒off where photosynthate is diverted from above‒ground to below-ground. As a non-photosynthetic organ, the plant root system requires photosynthates produced by shoots to maintain its metabolism, of which demand increases with root age (Eissenstat & Volder, 2005). The carbon cost of a root system can be substantial, accounting for as much as > 50% of the total produced photosynthates (Lambers, Atkin, & Millenaar, 2002). Particularly under water‒ or nutrient‒ limited conditions, plants tend to increase root-to-shoot ratio by directing more biomass to roots to enhance soil exploration (Sharp et al., 2003; Lopez et al., 2023). If plants can access the resources they need through a more efficient root system rather than a larger one, this would have significant benefits for overall plant health and productivity.

Our study showed that the narrow genotypes possessed compact root systems, whereas the wide genotypes possessed laterally spread root systems. This supported a consistent direction of root development from seedling to late-tillering growth stages. In line with previous research that has shown a good correlation between SRA and the root angle of mature plants in the field (Maccaferri et al., 2016; Alahmad et al., 2019), it implies that there is some degree of common genetic control of root angle across developmental stages, potentially enabling the use of high-throughput measurements in controlled environments to predict or select for adaptation in the field. Similar relationships between the angle of root axes and spatial patterns of root distribution have also been reported in other crops such as bread wheat and rice (Kato, Abe, Kamoshita, & Yamagishi, 2006; Manschadi et al., 2006; Manschadi, Hammer, Christopher, & deVoil, 2008).

### Potential implications of selection for seminal root angle on the architecture and functioning of the root system

In addition to root growth direction, the investigation of RSA based on the multiple factor analysis also revealed differences between wide and narrow genotypes in other root morphology traits. For example, the wide genotypes had a greater number of thinner roots, whereas the narrow genotypes tended to have fewer and thicker roots. This suggests the potential trade-off among some root morphology traits within the root system (i.e., root number and root diameter), as confirmed by RV coefficients that highlighted multiple covariations between many root trait groups.

Such trade-offs can be caused by physiological and/or genetic constraints (Weih, 2003). On the one hand, changes in RSA in relation to SRA did not influence the root biomass accumulation in the rhizobox study. This indicates a root morphological trade-off at the physiological level. The expression of root traits has a carbon cost. Therefore, for a given biomass investment in root systems, the promotion of certain root traits may come at the expense of other root traits. So it is important to determine the generality of our observations, understanding to what extent SRA can influence the inter-connections among different root morphological traits. On the other hand, the correlated response of root morphological traits to the selection for SRA could arise from the shared genetic control and/or linkage disequilibrium among different QTL. Previous studies have found some SRA QTL in durum wheat with pleiotropic effects controlling several root morphological traits (Maccaferri et al., 2016; Alemu et al., 2021). This suggested that root morphological trade-off might be genetically determined, and SRA might have an inherent effect on it. This being said, selecting for SRA may simultaneously select for some root morphological traits in a certain direction. For instance, selection for narrow SRA may favour selection for thick roots and small number of roots. As such, from a breeding perspective, the manipulation of SRA may constrain the genetic improvement of other root traits. Using SRA to identify optimal root systems would require a good understanding of the physiological and genetic links between SRA and other root traits. This knowledge will enhance our capacity to selectively breed for desired RSA.

Root traits are coordinated to shape the function of the root system. Their different expression reflects different resource acquisition strategies. The root traits observed in the wide-angle genotypes, such as a greater number of roots and roots with shallow angles, favour intensive soil exploration particularly for local resources e.g., water and nutrients. Small-diameter roots in the wide-angle genotypes also indicate high ratios of surface area to volume of roots, which increases contact with the soil, thereby enhancing the efficiency in soil resource capture. In contrast, narrow-angle genotypes exhibit root traits that are better suited for mining soil resources. Steep root angle combined with large-diameter roots increases the root penetration into deeper soil layers (Clark, Price, Steele, & Whalley, 2008; Lynch et al., 2021), enhancing the access to mobile soil resources, like water and nitrogen. Furthermore, larger root diameter has been linked to higher root hydraulic conductance that can improve water uptake and transport (Wan, Xu, Sosebee, Machado, & Archer, 2000). These variations in RSA suggest that SRA may be a key attribute that helps to identify genotypes with a RSA better adapted to target production environments. Narrow SRA could contribute to subsoil exploitation and provide advantages in environments where crops largely rely on deep water reserves, especially in terminal drought scenarios. On the other hand, wide SRA could support topsoil foraging, delivering benefits in environments characterised by limited availability of relatively immobile nutrients like phosphorus, or capturing in-season rainfall that has not yet leached to the subsoil.

### Outlook for exploring seminal root angle in future research

The results presented here suggest that narrow SRA genotypes have the propensity to develop deeper root systems at later stages of development. However, the differences in the RSAs between the wide and narrow genotypes were not statistically significant in the 60 ‒ 90 cm soil section (Figure 5). This could be due to several reasons. For instance, roots in most rhizoboxes were found to have already occupied the bottom section of the rhizobox when imaged. Hence, the time at which the experiment ceased could have been too late to observe the difference in root systems in deep soil. Moreover, we noted that although there is a general difference in RSA between the wide and narrow genotypes, for a specific genotype, its RSA may not match its SRA phenotype. For example, the NAM line 1_99 is a wide genotype but its ranking for PC1 at different depths implies that its RSA resembles that of the narrow genotypes (**Error! Reference source not found.**). Therefore, factors other than the genetic loci controlling SRA might also have influenced the phenotypic expression of genetically pre-defined RSA. Hence, to better understand how SRA affects RSA, future research will need to account for genetic background, whose dissimilarities have the potential to contribute to variations in root phenotypes that might obscure the impact of SRA.

The rhizobox experiment conducted in this study measured root systems of plants after six weeks of growth at the late-tillering stage. The stability of phenotype expression during development is an important criterion in plant breeding when selecting for a trait assessed at very early growth stages. Our study indicates that SRA allows some inferences to be made about RSA in older plants, which suggests that its association with the architecture of fully matured root systems merits investigation. Moreover, the current study conducted root phenotyping under controlled conditions that limit environmental influence. Given the plasticity of roots around competing acquisition of soil resources, conducting studies in controlled environments with limited water and/or nutrients will provide insights into root ideotypes for breeding more stress-resilient and resource-efficient crops. Lastly, since the ultimate benefit of high-throughput phenotypic screening is dependent on validation of useful root traits in the field for selection, it is also crucial to understand the expression of such root phenotypes in response to field environments.

## Conclusion

In this study, a set of genotypes with contrasting seedling SRAs were examined for their RSAs under controlled conditions. The significant differences in RSA between the wide and narrow genotypes demonstrate the prolonged effect of SRA on root development in the plants during later stages, which is likely independent of root:shoot biomass partitioning. Wide SRA showed a topsoil foraging root system that results from shallow angles of roots and high density/length of thin roots. In contrast, the RSA of narrow SRA favours deeper rooting, due to steep angles of roots and high proportion of thick roots. The results presented here support SRA of seedlings as a promising target for selection in breeding programs that aim to optimise root systems for improved crop resilience. Nevertheless, RSA is a complex trait, of which genetic variation is not only mediated by SRA. Therefore, further research into germplasm with more uniform genetic backgrounds to better evaluate SRA, particularly on fully mature root systems in the field, is required.

## Statements and declarations

## Acknowledgement

We are grateful for technical support provided by Dr Nora Derbal.

## Author contribution

YK, SA and LTH conceived and designed the study. YK performed the phenotyping. YK analysed the data. DRJ advised on the data analysis. YK wrote the manuscript. All authors read and reviewed the manuscript.

## Data availability

All data supporting the findings of this study are available in supplemental materials.

## References

Abdi, H., Williams, L. J., & Valentin, D. (2013). Multiple factor analysis: principal component analysis for multitable and multiblock data sets. WIREs Computational Statistics, 5(2), 149–179. 10.1002/wics.1246

Adiku, S. G. K., Ozier-Lafontaine, H., & Bajazet, T. (2001). Patterns of root growth and water uptake of a maize-cowpea mixture grown under greenhouse conditions. Plant and Soil, 235(1), 85–94. doi:10.1023/a:1011847214706

Adu, M. O., Asare, P. A., Yawson, D. O., Amoah, K. K., Atiah, K., Duah, M. K., & Graham, A. (2022). Root system traits contribute to variability and plasticity in response to phosphorus fertilization in 2 field-grown sorghum [Sorghum bicolor (L.) Moench] cultivars. Plant Phenomics. doi:doi:10.34133/plantphenomics.0002

Alahmad, S., El Hassouni, K., Bassi, F. M., Dinglasan, E., Youssef, C., Quarry, G., … Hickey, L. T. (2019). A major root architecture QTL responding to water limitation in durum wheat. Frontiers in Plant Science. doi:doi:10.3389/fpls.2019.00436

Alahmad, S., Kang, Y. C., Dinglasan, E., Jambuthenne, D., Robinson, H., Tao, Y. F., … Hickey, L. T. (2022). A multi-reference parent nested-association mapping population to dissect the genetics of quantitative traits in durum wheat. Genetic Resources and Crop Evolution. doi:doi:10.1007/s10722-022-01515-2

Alemu, A., Feyissa, T., Maccaferri, M., Sciara, G., Tuberosa, R., Ammar, K., … Abeyo, B. (2021). Genome-wide association analysis unveils novel QTLs for seminal root system architecture traits in Ethiopian durum wheat. Bmc Genomics. doi:doi:10.1186/s12864-020-07320-4

Aziz, M. M., Palta, J. A., Siddique, K. H. M., & Sadras, V. O. (2017). Five decades of selection for yield reduced root length density and increased nitrogen uptake per unit root length in Australian wheat varieties. Plant and Soil, 413(1-2), 181–192. doi:10.1007/s11104-016-3059-y

Borrell, A. K., Mullet, J. E., George-Jaeggli, B., van Oosterom, E. J., Hammer, G. L., Klein, P. E., & Jordan, D. R. (2014). Drought adaptation of stay-green sorghum is associated with canopy development, leaf anatomy, root growth, and water uptake. Journal of Experimental Botany, 65(21), 6251–6263. doi:10.1093/jxb/eru232

Borrell, A. K., Wong, A. C. S., George-Jaeggli, B., van Oosterom, E. J., Mace, E. S., Godwin, I. D., … Jordan, D. R. (2022). Genetic modification of PIN genes induces causal mechanisms of stay-green drought adaptation phenotype. Journal of Experimental Botany, 73(19), 6711–6726. doi:10.1093/jxb/erac336

Cane, M. A., Maccaferri, M., Nazemi, G., Salvi, S., Francia, R., Colalongo, C., & Tuberosa, R. (2014). Association mapping for root architectural traits in durum wheat seedlings as related to agronomic performance. Molecular Breeding, 34(4), 1629–1645. doi:10.1007/s11032-014-0177-1

Christopher, M., Chenu, K., Jennings, R., Fletcher, S., Butler, D., Borrell, A., & Christopher, J. (2018). QTL for stay-green traits in wheat in well-watered and water-limited environments. Field Crops Research, 217, 32–44. doi:10.1016/j.fcr.2017.11.003

Clark, L. J., Price, A. H., Steele, K. A., & Whalley, W. R. (2008). Evidence from near-isogenic lines that root penetration increases with root diameter and bending stiffness in rice. Functional Plant Biology, 35(11), 1163–1171. doi:10.1071/fp08132

Eissenstat, D. M., & Volder, A. (2005). The efficiency of nutrient acquisition over the life of a root. In Nutrient acquisition by plants: an ecological perspective (pp. 185–220). Berlin, Heidelberg: Springer Berlin Heidelberg.

Figueroa-Bustos, V., Palta, J. A., Chen, Y. L., & Siddique, K. H. (2018). Characterization of root and shoot traits in wheat cultivars with putative differences in root system size. Agronomy-Basel. doi:doi:10.3390/agronomy8070109

Figueroa-Bustos, V., Palta, J. A., Chen, Y. L., Stefanova, K., & Siddique, K. H. M. (2020). Wheat cultivars with contrasting root system size responded differently to terminal drought. Frontiers in Plant Science. doi:doi:10.3389/fpls.2020.01285

Johnson, M. G., Tingey, D. T., Phillips, D. L., & Storm, M. J. (2001). Advancing fine root research with minirhizotrons. Environmental and Experimental Botany, 45(3), 263–289. doi:10.1016/s0098-8472(01)00077-6

Kato, Y., Abe, J., Kamoshita, A., & Yamagishi, J. (2006). Genotypic variation in root growth angle in rice (Oryza sativa L.) and its association with deep root development in upland fields with different water regimes. Plant and Soil, 287(1-2), 117–129. doi:10.1007/s11104-006-9008-4

Lambers, H., Atkin, O. K., & Millenaar, F. F. (2002). Respiratory patterns in roots in relation to their functioning. In Plant roots: the hidden half (3rd edn ed., pp. 521–552). New York, NY, USA: Marcel Dekker.

Le, S., Josse, J., & Husson, F. (2008). FactoMineR: An R package for multivariate analysis. Journal of Statistical Software, 25(1), 1–18. doi:10.18637/jss.v025.i01

Lopez, G., Ahmadi, S. H., Amelung, W., Athmann, M., Ewert, F., Gaiser, T., … Seidel, S. J. (2023). Nutrient deficiency effects on root architecture and root-to-shoot ratio in arable crops. Frontiers in Plant Science. doi:doi:10.3389/fpls.2022.1067498

Lynch, J. P. (2007). Roots of the second green revolution. Australian Journal of Botany, 55(5), 493–512. doi:10.1071/bt06118

Lynch, J. P., Strock, C. F., Schneider, H. M., Sidhu, J. S., Ajmera, I., Galindo-Castaneda, T., … Hanlon, M. T. (2021). Root anatomy and soil resource capture. Plant and Soil, 466(1-2), 21–63. doi:10.1007/s11104-021-05010-y

Maccaferri, M., El-Feki, W., Nazemi, G., Salvi, S., Cane, M. A., Colalongo, M. C., … Tuberosa, R. (2016). Prioritizing quantitative trait loci for root system architecture in tetraploid wheat. Journal of Experimental Botany, 67(4), 1161–1178. doi:10.1093/jxb/erw039

Mace, E. S., Singh, V., Van Oosterom, E. J., Hammer, G. L., Hunt, C. H., & Jordan, D. R. (2012). QTL for nodal root angle in sorghum (Sorghum bicolor L. Moench) co-locate with QTL for traits associated with drought adaptation. Theoretical and Applied Genetics, 124(1), 97–109. doi:10.1007/s00122-011-1690-9

Manschadi, A. M., Christopher, J., Devoil, P., & Hammer, G. L. (2006). The role of root architectural traits in adaptation of wheat to water-limited environments. Functional Plant Biology, 33(9), 823–837. doi:10.1071/fp06055

Manschadi, A. M., Hammer, G. L., Christopher, J. T., & deVoil, P. (2008). Genotypic variation in seedling root architectural traits and implications for drought adaptation in wheat (Triticum aestivum L.). Plant and Soil, 303(1-2), 115–129. doi:10.1007/s11104-007-9492-1

Menamo, T., Borrell, A. K., Mace, E., Jordan, D. R., Tao, Y. F., Hunt, C., & Kassahun, B. (2023). Genetic dissection of root architecture in Ethiopian sorghum landraces. Theoretical and Applied Genetics, 136(10). doi:10.1007/s00122-023-04457-0

Nagel, K. A., Putz, A., Gilmer, F., Heinz, K., Fischbach, A., Pfeifer, J., … Schurr, U. (2012). GROWSCREEN-Rhizo is a novel phenotyping robot enabling simultaneous measurements of root and shoot growth for plants grown in soil-filled rhizotrons. Functional Plant Biology, 39(10-11), 891–904. doi:10.1071/fp12023

Ober, E. S., Alahmad, S., Cockram, J., Forestan, C., Hickey, L. T., Kant, J., … Watt, M. (2021). Wheat root systems as a breeding target for climate resilience. Theoretical and Applied Genetics, 134(6), 1645–1662. doi:10.1007/s00122-021-03819-w

Palta, J. A., Chen, X., Milroy, S. P., Rebetzke, G. J., Dreccer, M. F., & Watt, M. (2011). Large root systems: are they useful in adapting wheat to dry environments? Functional Plant Biology, 38(5), 347–354. doi:10.1071/fp11031

R Core Team. (2013). R: A language and environment for statistical computing. Vienna, Austria: R Foundation for Statistical Computing.

Rich, S. M., & Watt, M. (2013). Soil conditions and cereal root system architecture: review and considerations for linking Darwin and Weaver. Journal of Experimental Botany, 64(5), 1193–1208. doi:doi:10.1093/jxb/ert043

Richard, C. A. I., Hickey, L. T., Fletcher, S., Jennings, R., Chenu, K., & Christopher, J. T. (2015). High-throughput phenotyping of seminal root traits in wheat. Plant Methods. doi:doi:10.1186/s13007-015-0055-9

Robinson, H., Hickey, L., Richard, C., Mace, E., Kelly, A., Borrell, A., … Fox, G. (2016). Genomic Regions Influencing Seminal Root Traits in Barley. Plant Genome, 9(1). doi:10.3835/plantgenome2015.03.0012

Seethepalli, A., Dhakal, K., Griffiths, M., Guo, H., Freschet, G. T., & York, L. M. (2021). RhizoVision Explorer: Open-source software for root image analysis and measurement standardization. Aob Plants. doi:doi:10.1093/aobpla/plab056

Sharp, R. E., Bohnert, H. J., Springer, G. K., Davis, G. L., Schachtman, D. P., Wu, Y. J., & Nguyen, H. T. (2003). Root growth maintenance during water deficits: physiology to functional genomics. Journal of Experimental Botany, 54, 20–20. Retrieved from<GO to ISI>://WOS:000224244600063

Singh, V., van Oosterom, E. J., Jordan, D. R., & Hammer, G. L. (2012). Genetic control of nodal root angle in sorghum and its implications on water extraction. European Journal of Agronomy, 42, 3–10. doi:10.1016/j.eja.2012.04.006

Trachsel, S., Kaeppler, S. M., Brown, K. M., & Lynch, J. P. (2011). Shovelomics: high throughput phenotyping of maize (Zea mays L.) root architecture in the field. Plant and Soil, 341(1-2), 75–87. doi:10.1007/s11104-010-0623-8

van Oosterom, E. J., Yang, Z. J., Zhang, F. L., Deifel, K. S., Cooper, M., Messina, C. D., & Hammer, G. L. (2016). Hybrid variation for root system efficiency in maize: potential links to drought adaptation. Functional Plant Biology, 43(6), 502–511. doi:10.1071/fp15308

Waines, J. G., & Ehdaie, B. (2007). Domestication and crop physiology: Roots of green-revolution wheat. Annals of Botany, 100(5), 991–998. doi:10.1093/aob/mcm180

Wan, C. G., Xu, W. W., Sosebee, R. E., Machado, S., & Archer, T. (2000). Hydraulic lift in drought-tolerant and -susceptible maize hybrids. Plant and Soil, 219(1-2), 117–126. doi:10.1023/a:1004740511326

Wasson, A. P., Rebetzke, G. J., Kirkegaard, J. A., Christopher, J., Richards, R. A., & Watt, M. (2014). Soil coring at multiple field environments can directly quantify variation in deep root traits to select wheat genotypes for breeding. Journal of Experimental Botany, 65(21), 6231–6249. doi:10.1093/jxb/eru250

Weih, M. (2003). Trade-offs in plants and the prospects for breeding using modern biotechnology. New Phytologist, 158(1), 7–9. doi:10.1046/j.1469-8137.2003.00716.x

Zhang, L., Du, Y. L., & Li, X. G. (2020). Modern wheat cultivars have greater root nitrogen uptake efficiency than old cultivars. Journal of Plant Nutrition and Soil Science, 183(2), 192–199. doi:10.1002/jpln.201900353

