## Supplemental materials for "Seminal root angle is associated with root system architecture in durum wheat"

**Supplemental Tables**

Supplemental Table S1. Proportional contribution of root groups to the first two principal components (PC1 and PC2) of the multiple factor analysis conducted on the intact root system (0 – 90 cm) as well as on the roots in different soil sections (0 ‒ 30, 30 ‒ 60, and 60 – 90 cm).

|  |  | **Contribution (%)** | |
| --- | --- | --- | --- |
| **Multiple factor analysis** | **Root trait group** | **PC1** | **PC2** |
| 0 ‒ 30 cm | Angle | 9.4 | 49.0 |
|  | Area | 17.8 | 15.1 |
|  | Diameter | 18.1 | 0.9 |
|  | Length | 17.1 | 13.1 |
|  | Number | 18.4 | 20.5 |
|  | Volume | 19.1 | 1.5 |
| 30 ‒ 60 cm | Angle | 10.7 | 23.7 |
|  | Area | 17.8 | 31.5 |
|  | Diameter | 18.3 | 0.4 |
|  | Length | 15.3 | 19.1 |
|  | Number | 19.0 | 17.8 |
|  | Volume | 18.9 | 7.6 |
| 60 ‒ 90 cm | Angle | 12.4 | 32.7 |
|  | Area | 17.5 | 26.1 |
|  | Diameter | 16.7 | 7.4 |
|  | Length | 17.3 | 13.3 |
|  | Number | 18.2 | 17.0 |
|  | Volume | 18.0 | 3.6 |
| Overall (0 ‒ 90 cm) | Angle | 11.2 | 25.7 |
|  | Area | 17.3 | 32.8 |
|  | Diameter | 18.0 | 0.7 |
|  | Length | 16.3 | 17.5 |
|  | Number | 18.6 | 19.6 |
|  | Volume | 18.6 | 3.7 |

Supplemental Table S2. Genotype means for the scores of the first two PCs of each multiple factor analysis.

| **Multiple factor analysis** | **Genotype** | **Group** | **PC1** | **PC2** |
| --- | --- | --- | --- | --- |
| 0 ‒ 30 cm | 1_107 | Wide | 0.89 | -0.43 |
|  | 1_17 | Wide | 0.31 | 0.27 |
|  | 1_42 | Wide | 0.55 | -0.75 |
|  | 1_63 | Wide | 0.41 | -0.31 |
|  | 1_99 | Wide | -0.45 | -0.38 |
|  | 3_51 | Narrow | -1.66 | 0.49 |
|  | 3_86 | Narrow | -1.13 | 0.6 |
|  | 3_88 | Narrow | -0.08 | 0.46 |
|  | 6_17 | Narrow | -1.99 | 0.5 |
|  | 6_21 | Narrow | 0.09 | -0.15 |
|  | DBA Aurora | Wide | 3.06 | -0.29 |
| 30 ‒ 60 cm | 1_107 | Wide | 1.9 | 0.15 |
|  | 1_17 | Wide | 0.12 | 0.26 |
|  | 1_42 | Wide | 0.13 | -0.91 |
|  | 1_63 | Wide | -0.26 | -0.66 |
|  | 1_99 | Wide | -0.37 | 0.01 |
|  | 3_51 | Narrow | -0.15 | 0.79 |
|  | 3_86 | Narrow | 0.41 | 0.11 |
|  | 3_88 | Narrow | 0.39 | -0.25 |
|  | 6_17 | Narrow | -1.53 | 0 |
|  | 6_21 | Narrow | -1.38 | 0.37 |
|  | DBA Aurora | Wide | 0.73 | 0.14 |
| 60 ‒ 90 cm | 1_107 | Wide | 1.68 | -0.49 |
|  | 1_17 | Wide | 0.55 | 0.5 |
|  | 1_42 | Wide | -0.17 | -0.19 |
|  | 1_63 | Wide | 0.61 | -0.57 |
|  | 1_99 | Wide | -1.02 | 0.04 |
|  | 3_51 | Narrow | 0.17 | 0.65 |
|  | 3_86 | Narrow | 0.96 | 0.22 |
|  | 3_88 | Narrow | -0.98 | -0.34 |
|  | 6_17 | Narrow | -0.87 | 0.05 |
|  | 6_21 | Narrow | 0.14 | 0.14 |
|  | DBA Aurora | Wide | -1.06 | -0.01 |
| Overall (0 ‒ 90 cm) | 1_107 | Wide | 2.14 | -0.37 |
|  | 1_17 | Wide | 0.54 | 0.55 |
|  | 1_42 | Wide | 0.15 | -0.75 |
|  | 1_63 | Wide | 0.34 | -0.61 |
|  | 1_99 | Wide | -0.82 | -0.06 |
|  | 3_51 | Narrow | -0.72 | 0.62 |
|  | 3_86 | Narrow | 0.13 | 0.15 |
|  | 3_88 | Narrow | -0.45 | -0.13 |
|  | 6_17 | Narrow | -1.94 | 0.11 |
|  | 6_21 | Narrow | -0.54 | 0.28 |
|  | DBA Aurora | Wide | 1.18 | 0.22 |

Supplemental Table S3. Comparison between wide and narrow seminal root angle genotypes for PC1 and PC2 of each multiple factor analysis using the *t*-test or post-hoc test.

| **Multiple factor analysis** | **PC** | **Mean of group** | | ***P*-value** |
| --- | --- | --- | --- | --- |
|  |  | **Wide** | **Narrow** |  |
| 0 ‒ 30 cm | PC1 | 0.79 | -0.95 | < 0.01 |
|  | PC2 | -0.32 | 0.38 | < 0.01 |
| 30 ‒ 60 cm | PC1 | 0.38 | -0.45 | 0.14 |
|  | PC2 | -0.17 | 0.21 | 0.21 |
| 60 ‒ 90 cm | PC1 | 0.10 | -0.12 | 0.71 |
|  | PC2 | -0.12 | 0.14 | 0.23 |
| Overall (0 ‒ 90 cm) | PC1 | 0.59 | -0.7 | < 0.05 |
|  | PC2 | -0.17 | 0.2 | 0.16 |

Supplemental Table S4. The results of the Pearson’s correlation analysis between the PC scores of the multiple factor analysis conducted on the intact root system (0 – 90 cm) and on the roots in the 0 ‒ 30 cm soil section, and the root imaging traits that showed a significant correlation with the PCs, including their significance and correlation coefficients.

| **Multiple factor analysis** | **PC** | ***P‒*value** | **Coeff.** | **Trait** | **Root trait group** |
| --- | --- | --- | --- | --- | --- |
| 0 ‒ 30 cm | PC1 | 2.87E-10 | -0.69 | ARO | Angle |
|  |  | 5.7E-40 | 0.97 | PAD2 | Area |
|  |  | 5.81E-40 | 0.97 | SAD2 | Area |
|  |  | 1.18E-34 | 0.96 | PAD1 | Area |
|  |  | 1.24E-34 | 0.96 | SAD1 | Area |
|  |  | 2.05E-18 | -0.85 | Solidity | Area |
|  |  | 3.29E-16 | -0.82 | PAD3 | Area |
|  |  | 3.69E-16 | -0.82 | SAD3 | Area |
|  |  | 1.91E-13 | -0.77 | SA | Area |
|  |  | 2.85E-06 | -0.55 | NeA | Area |
|  |  | 9.05E-05 | -0.47 | AHS | Area |
|  |  | < 0.05 | -0.31 | LRA | Area |
|  |  | < 0.05 | -0.29 | CoA | Area |
|  |  | 9.19E-40 | -0.97 | AvD | Diameter |
|  |  | 5.28E-34 | -0.96 | MeD | Diameter |
|  |  | 2.65E-09 | -0.67 | MxD | Diameter |
|  |  | 4.35E-40 | 0.97 | RLD2 | Length |
|  |  | 3.05E-37 | 0.97 | Perimeter | Length |
|  |  | 5.04E-35 | 0.96 | RLD1 | Length |
|  |  | 5.34E-19 | 0.85 | TRL | Length |
|  |  | 1.98E-10 | 0.70 | RLD3 | Length |
|  |  | 1.15E-25 | 0.91 | MNR | Number |
|  |  | 3.08E-25 | 0.91 | MxNR | Number |
|  |  | 1.23E-24 | 0.91 | NRT | Number |
|  |  | 9.4E-23 | 0.89 | Holes | Number |
|  |  | 5.71E-13 | 0.76 | ShAF | Number |
|  |  | 2.16E-07 | -0.60 | MAF | Number |
|  |  | 8.47E-40 | 0.97 | VDR2 | Volume |
|  |  | 1.24E-34 | 0.96 | VDR1 | Volume |
|  |  | 6.15E-29 | -0.93 | VDR3 | Volume |
|  |  | 8.68E-29 | -0.93 | Volume | Volume |
|  | PC2 | 2.28E-09 | 0.67 | ARO | Angle |
|  |  | 1.13E-05 | 0.52 | NeA | Area |
|  |  | 2.10E-08 | 0.64 | CoA | Area |
|  |  | < 0.05 | 0.32 | SA | Area |
|  |  | < 0.05 | 0.28 | PAD3 | Area |
|  |  | < 0.05 | 0.28 | SAD3 | Area |
|  |  | < 0.001 | 0.41 | TRL | Length |
|  |  | 4.44E-06 | 0.54 | RLD3 | Length |
|  |  | < 0.05 | 0.28 | Holes | Number |
|  |  | < 0.01 | -0.38 | ShAF | Number |
|  |  | 1.52E-10 | 0.70 | StAF | Number |
| Overall (0 ‒ 90 cm) | PC1 | 3.92E-13 | -0.76 | ARO | Angle |
|  |  | 5.94E-35 | 0.96 | PAD2 | Area |
|  |  | 6.21E-35 | 0.96 | SAD2 | Area |
|  |  | 2.83E-31 | 0.94 | PAD1 | Area |
|  |  | 3.07E-31 | 0.94 | SAD1 | Area |
|  |  | 2.41E-24 | -0.91 | Solidity | Area |
|  |  | 5.24E-10 | -0.69 | PAD3 | Area |
|  |  | 5.66E-10 | -0.69 | SAD3 | Area |
|  |  | 6.81E-08 | -0.62 | SA | Area |
|  |  | < 0.01 | -0.37 | NeA | Area |
|  |  | 8.83E-41 | -0.97 | AvD | Diameter |
|  |  | 2.62E-35 | -0.96 | MeD | Diameter |
|  |  | < 0.01 | -0.38 | MxD | Diameter |
|  |  | 7.07E-37 | 0.96 | Perimeter | Length |
|  |  | 3.46E-35 | 0.96 | RLD2 | Length |
|  |  | 2.50E-31 | 0.95 | RLD1 | Length |
|  |  | 3.03E-16 | 0.82 | TRL | Length |
|  |  | 2.96E-09 | 0.66 | RLD3 | Length |
|  |  | 1.49E-25 | 0.91 | NRT | Number |
|  |  | 2.40E-19 | 0.86 | Holes | Number |
|  |  | 4.94E-17 | 0.83 | MNR | Number |
|  |  | 9.08E-13 | 0.75 | ShAF | Number |
|  |  | 3.49E-10 | 0.69 | MxNR | Number |
|  |  | 1.04E-08 | -0.65 | MAF | Number |
|  |  | < 0.001 | -0.43 | StAF | Number |
|  |  | 1.15E-34 | 0.96 | VDR2 | Volume |
|  |  | 3.07E-31 | 0.94 | VDR1 | Volume |
|  |  | 9.28E-23 | -0.89 | VDR3 | Volume |
|  |  | 1.25E-22 | -0.89 | Volume | Volume |
